# A novel reticular oscillator in the brainstem synchronizes neonatal crying with breathing

**DOI:** 10.1101/2021.02.26.433060

**Authors:** Xin Paul Wei, Matthew Collie, Bowen Dempsey, Gilles Fortin, Kevin Yackle

**Affiliations:** Department of Physiology, University of California-San Francisco, San Francisco, CA 94143, USA; Biomedical Sciences Graduate Program, University of California-San Francisco, San Francisco, CA 94143, USA; Institut de Biologie de l’École Normale Supérieure (IBENS), École Normale Supérieure, CNRS, INSERM, PSL Research University, Paris, France

**Author notes:** Program in Neuroscience, Harvard University, Boston, MA 02115, USA.

## Abstract

Human speech can be divided into short, rhythmically-timed elements, similar to syllables within words. Even our cries and laughs, as well as the vocalizations of other species, are periodic. However, the cellular and molecular mechanisms underlying the tempo of mammalian vocalizations remain unknown. Here we describe rhythmically-timed neonatal mouse vocalizations that occur within single breaths, and identify a brainstem node that structures these cries, which we name the intermediate reticular oscillator (iRO). We show that the iRO acts autonomously and sends direct inputs to key muscles in order to coordinate neonatal vocalizations with breathing, as well as paces and patterns these cries. These results reveal that a novel mammalian brainstem oscillator embedded within the conserved breathing circuitry plays a central role in the production of neonatal vocalizations.

## Main Text

Rhythmicity underlies both human speech and animal vocalizations. For instance, speech oscillates in volume with each syllable (*1*), the communicative calls of marmosets are composed of repeating units (*2*), and the timing of songbird syllables are regularly spaced as they learn to sing (*3*). These and other examples suggest that the tempo of sounds within vocalizations is innately encoded. Indeed, the rhythmic vocalizations of midshipman fish are timed by pacemaker neurons in the hindbrain (*4*). It has therefore been hypothesized that the rhythmicity of mammalian sound production is created by hardwired neural circuits in the brainstem, but evidence to support this theory is lacking.

The innate vocalization motor program is initiated by brainstem projections from the midbrain periaqueductal gray (*5*); however, the neural circuitry required to fully orchestrate these vocalizations remains poorly understood (*6*, *7*). Moreover, because mammalian vocalizations are produced by the concerted activity of articulator (laryngeal and tongue) and breathing (diaphragm and intercostal) muscles (*8*), they must be seamlessly integrated with the breathing rhythm. We hypothesized that the vocalization motor patterning system is anatomically and functionally connected to the neural circuit for breathing in the brainstem. We also predicted that this circuit encodes rhythmicity within vocalizations. Here, by studying the neural control of instinctive cries produced by neonatal mice, which are analogous to the cries of human infants (*9*–*11*), we elucidate the brainstem mechanism that patterns and paces mammalian vocalizations.

### Neonatal cries comprise syllables that are rhythmically timed within a breath

To study the coordination of vocalizations with breathing, we induced instinctive ultrasonic vocalizations (USVs) by removing neonatal mice from their nests. These cries were recorded while breathing was monitored by unrestrained barometric plethysmography (*12*) (Fig. 1A). Cries were characterized by increased inspiratory and expiratory airflow coupled with USVs in the 50-150 kHz range (Fig. 1A-C) and occurred in bouts of 9.7 ± 0.6 breaths (average ± SEM, n = 16 pups). Most cry breaths comprised one distinct vocalization, or syllable, that commenced ~10-20 milliseconds after the onset of expiration and coincided with peak expiratory airflow (Fig. 1D). However, cry breaths in the middle of a bout tended to include two, three, or sometimes more elements, and were therefore designated as multisyllabic (Fig. 1C-D). In these cases, the onset of a syllable was rhythmically timed to follow local peaks in expiratory airflow (Fig. 1D), and the syllables were separated by a period of silence associated with a dip in airflow. Syllables within unisyllabic cries were longer than those in multisyllabic cries, and syllables progressively increased in length during multisyllabic cries (Fig. 1E). The rhythmicity of postnatal mouse cries led us to hypothesize that they were timed by a fast conditional oscillator nested within a slower breathing pattern.

**Fig. 1.**
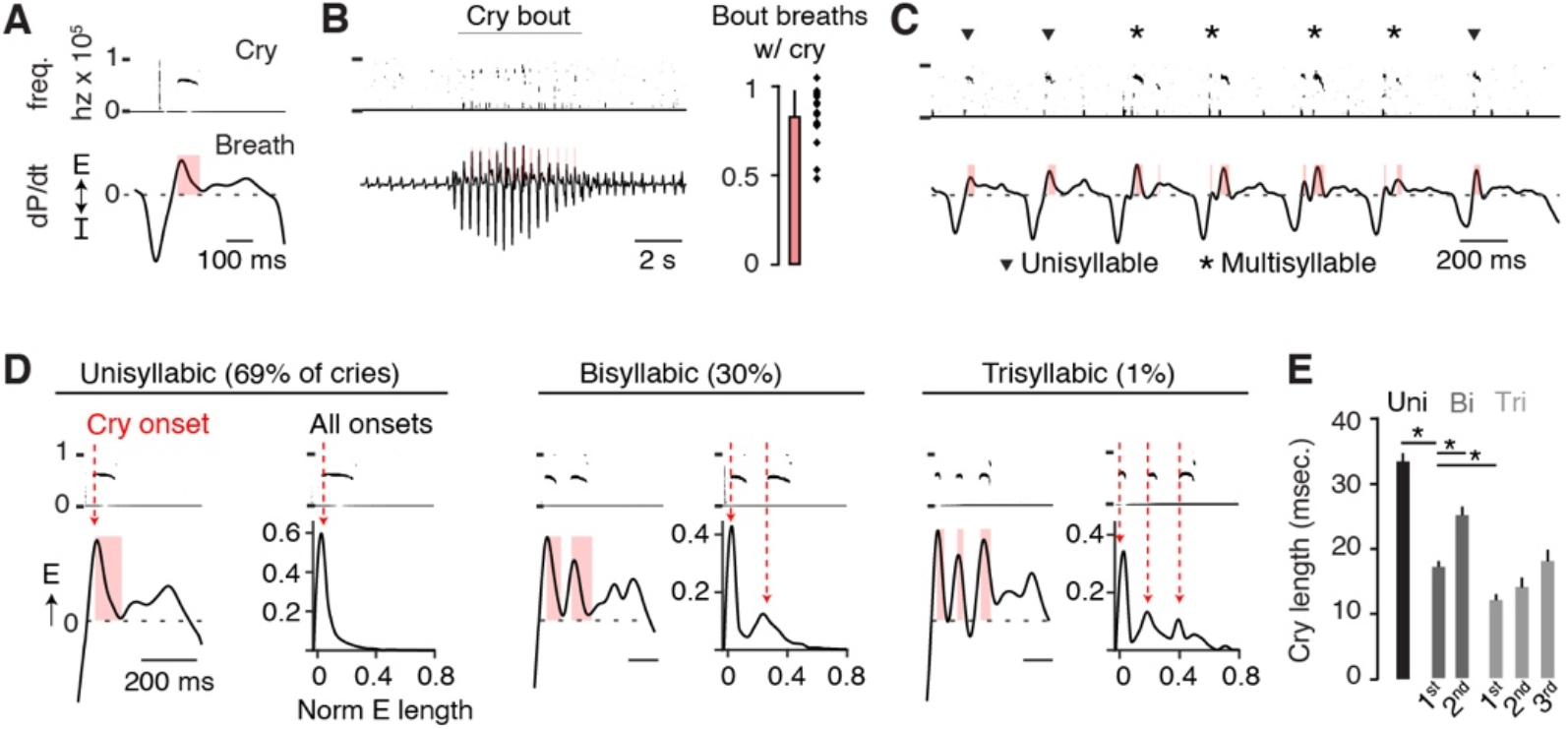
Neonatal cries comprise syllables that are rhythmically timed within a breath. **A**, Example cry breath with associated cry (frequency spectrogram 0-100 kHz) and respiratory pressure changes (dP/dt, arbitrary units). E and I, expiratory and inspiratory pressure change. Red bar indicates USV length. **B**, Example of large breaths during a cry bout and basal breaths before and after. Right, proportion of large cry bout breaths with cries (n = 17 mice, mean + SD) **C**, Enlargement cry bout in **B** showing four multisyllabic (*) and three unisyllabic (▾) cries. **D**, Left, representative examples of expiratory airflow when one, two, or three syllables are voiced within a single breath. Right, histograms of cry onset time normalized to expiratory duration (total n = 5596 cries from n = 23 mice). **E**, Syllable length within uni-, bi-, and trisyllabic cries. *, one-sided t-test p-value < 0.05.

### Cry syllables are synchronized with the respiratory motor program

Murine adult vocalizations are produced when airflow through a closed (adducted) glottis becomes rapid and then turbulent after colliding with the laryngeal cartilage (*13*, *14*). To explore whether neonatal cries similarly require laryngeal adduction, we disrupted the thyroarytenoid (TA) muscle, one of the key muscles that closes the larynx. Bilateral electrolytic lesion of the TA eliminated cries despite the retention of augmented airflow during cry bouts (Fig. 2A). Surprisingly, attempted cry breaths after TA lesion still showed airflow oscillations reminiscent of multisyllabic cries (Fig. 2B). This suggested that, although laryngeal adduction is required for producing the sound of each cry syllable, the airflow dip that separates the syllables is generated by a distinct mechanism.

**Fig. 2.**
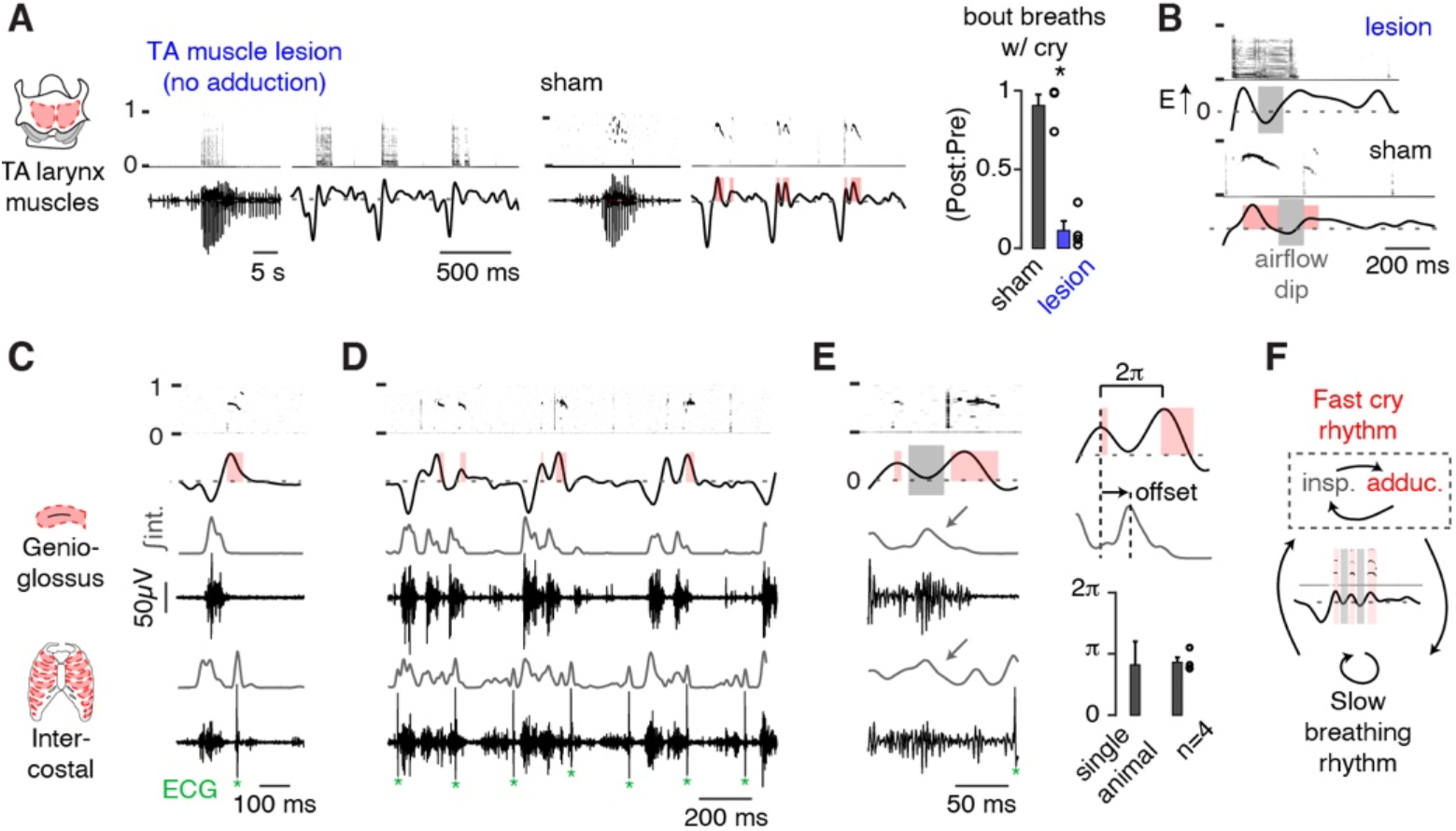
Cry syllables are synchronized with the respiratory motor program. **A**, Example cry bout and three cry breaths after bilateral electrolytic lesion of the TA laryngeal adductor muscle (left) and sham lesion (middle). Right, the number large cry bout breaths with cries in post-versus pre-surgery (post:pre) recordings. Sham, black, n = 3; TA lesion, red, n = 4; mean ± SEM. *, one-sided Wilcoxon rank sum p-value < 0.05. **B**, Cry breaths after TA lesion retain airflow oscillations reminiscent of multisyllabic cries in sham animals. **C**, Electromyographic (EMG) activity of the genioglossus (n = 4) and intercostal (n = 5) inspiratory muscles. Gray, integrated (∫int.) activity. Example unisyllabic cry and associated inspiratory muscle activity. * denotes contaminating cardiac ECG signals. **D**, Example multisyllabic calls with additional inspiratory activity during airflow dips. **E**, Left, example multisyllabic cry breath showing the gap between syllables coinciding with genioglossus and intercostal activity. Right, quantification of timing of genioglossus activity with respect to peak expiratory airflow for a single neonate and n = 4 animals. **F**, Schematic showing how cycling of inspiration and laryngeal adduction produces multisyllabic cries within a slower breathing rhythm.

Because the larynx adducts immediately after inspiration during basal breathing (*15*), we wondered if the airflow dips that preceded laryngeal adductions during cry syllables are also patterned by an inspiratory motor program. In this way, inspiration would be ectopically initiated during expiration. Indeed, signatures of the inspiratory motor program, such as genioglossus (tongue) and intercostal muscle activity (*16*), preceded each cry syllable during airflow dips (Fig. 2C-D). Inspiratory muscle activity was maximal approximately halfway between two expiratory airflow peaks (Fig. 2E), during which time, body wall movements mimicked inspiration (Fig. S1 and movie S1). These data suggest that the proposed oscillator is able to time multisyllable cries by cyclically re-engaging the respiratory motor program to generate inspiration followed by laryngeal adduction within a slower breath (Fig. 2F).

### The medullary brainstem contains a cluster of premotor neurons for sound articulation

To begin investigating the underlying circuit driving rhythmic cries, we first sought to identify premotor neurons to the key muscles that articulate sound, including the TA and cricothyroid (CT) muscles that adduct the larynx. We thus performed monosynaptic viral tracing from TA and CT motor neurons, using two methods: 1) microinjecting TA or CT muscles with glycoprotein-deleted rabies virus (ΔG-rabies-mCherry), complemented with glycoprotein-expressing herpes simplex virus (HSV-G) in wildtype neonates or 2) microinjecting these muscles with ΔG-rabies-mCherry in mice expressing the glycoprotein in all motor neurons (ChAT-Cre;ROSA-LSL-G-TVA) (Fig. 3A). We confirmed that laryngeal muscle injections were restricted to the intended muscle (Fig. S2). Four to seven days after injection, we found infected neurons arising from both muscles in similar areas throughout the ventral respiratory column (TA, n = 10; CT, n = 10) (Fig. 3A-B and Fig. S3). Traced premotor neurons were distinguished from motor neurons by the absence of Phox2b transcription factor expression (*17*) and were found medial to the compact nucleus ambiguus (cNA) in the rostral ventral intermediate reticular formation (rv-iRF), the Bötzinger complex (BötC), the preBötzinger complex (preBötC), and the retroambiguus (RAm) (Fig. 3A-B and Fig. S3-S4). To characterize the neurotransmitter used by these various TA premotor neuron pools, we injected ΔG-mCherry-rabies and HSV-G into the TA muscle of transgenic neonatal mice expressing labeled glutamatergic (Vglut2-Cre;ROSA-LSL-YFP) or GABAergic (Vgat-Cre;ROSA-LSL-YFP) neurons. TA premotor neurons within the rv-iRF were Vglut2-positive, those in other respiratory brainstem centers including the BötC and RAm were mostly Vgat-positive, and those in the preBötC were either Vglut2-positive or Vgat-positive (Fig. 3C-D).

**Fig. 3.**
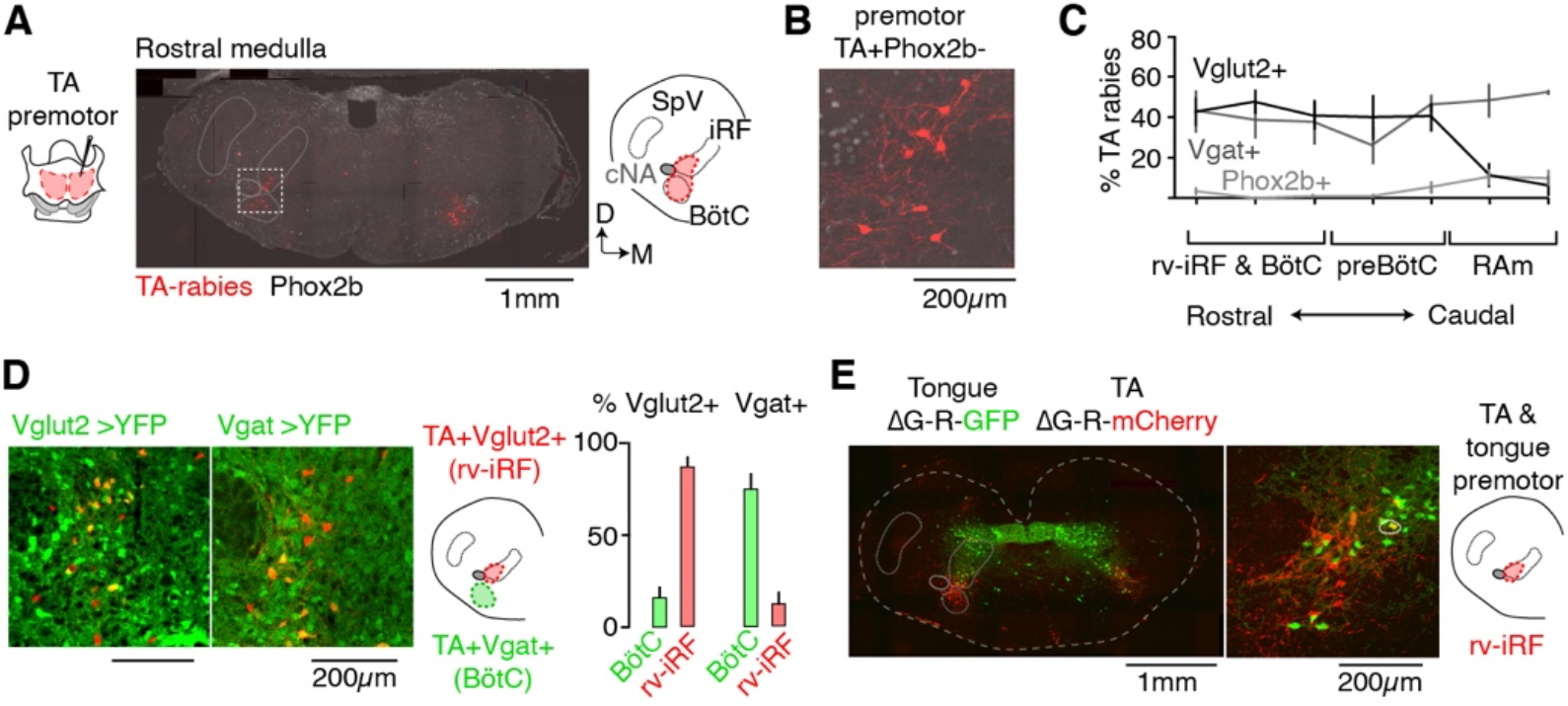
The medullary brainstem contains a cluster of premotor neurons for sound articulation. **A**, Monosynaptically-restricted rabies virus injected into the TA (ΔG-Rabies-mCherry and HSV-G). Middle, 20μm thick slice of the rostral medullary brainstem showing the distribution of rabies-mCherry-positive TA premotor neurons (red) and Phox2b expression (gray). Right, schematic of slice illustrating the position of the iRF and BötC, cNA, and spinal trigeminal nucleus (SpV). D, dorsal; M, medial. **B**, Left, magnification of boxed area in **A** showing a cluster of Phox2b-negative premotor TA-traced neurons directly medial to the cNA. **C**, Quantification of the mean percent ± SEM of rabies-infected neurons that were Vglut2-positive (black, n=3), Vgat-positive (gray, n=3), or Phox2B-positive (TA motor, pale gray, n=6) across the ventral respiratory centers (rv-iRF/BötC, preBötC, and RAm). **D**, Left, TA premotor neurons in the rv-iRF and BötC of Vglut2-Cre→YFP (n=3) and Vgat-Cre→YFP mice (n=3). Neurons medial to cNA are Vglut2-positive (rv-iRF) and those in ventral BötC are Vgat-positive. Right, quantification. **E**, Left, Slice of the rostral medullary brainstem showing the distribution of ΔG-rabies-mCherry-positive TA premotor neurons and ΔG-rabies-GFP-positive tongue premotor neurons. Middle, magnification of rv-iRF showing spatial overlap of TA and tongue premotor neurons. Circled neuron is double positive. Right, schematic of slice illustrating the position of the rv-iRF.

We next asked whether premotor neurons for other muscles of articulation, such as the tongue, overlap with one or more laryngeal premotor neuron clusters. To directly compare the spatial distribution of TA and tongue premotor neurons, we microinjected the tongue and TA with ΔG-rabies-expressing GFP or mCherry, respectively. Of the areas that contained TA premotor neurons, only the rv-iRF contained neurons premotor to the tongue (Fig. 3E). Importantly, some neurons in this region innervated both the TA and the tongue (Fig. 3E). Although the rv-iRF is distinct from brainstem sites that have previously been implicated in vocalization, such as the caudal RAm (*5*), it is apparently unique in containing excitatory premotor neurons for multiple cry articulators. It therefore became the prime candidate for our proposed oscillator to pace and pattern neonatal cries.

### The rv-iRF containing the premotor cluster is required for neonatal crying but not breathing

We sought to test the necessity of the rv-iRF for patterning neonatal crying by measuring breathing and cries before and after bilateral electrolytic lesion of the premotor cluster medial to the cNA. Each animal’s cry and breath parameters were measured 24 hours after lesion and normalized to pre-lesion data. The anatomical location of the lesions allowed us to classify them as bilateral on-target, bilateral off-target, or unilateral on-target (n = 5, 7 and 6, respectively). On-target lesions were limited to the region medial to the cNA, in the location containing the common cluster of cry articulator premotor neurons (Figs. 3 and 4B).

**Fig. 4.**
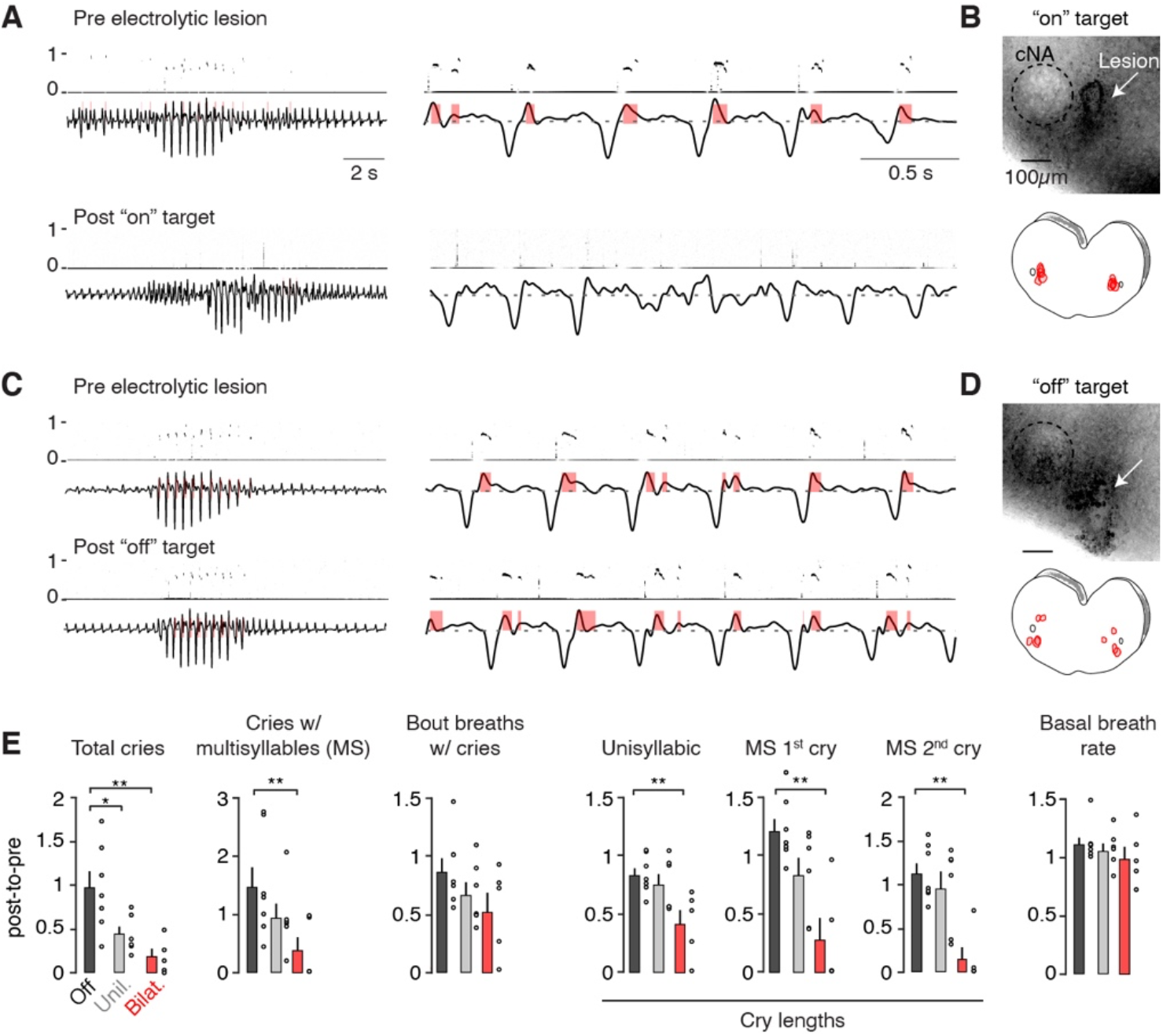
The rv-iRF premotor cluster is required for neonatal crying but not breathing. **A**, Representative cry bout and breathing pattern before (top) and after (bottom) bilateral on-target electrolytic lesion of the rv-iRF medial to cNA. **B**, Top, representative light microscopy image of the lesion. White arrow indicates lesion site. Bottom, summary of locations for all on-target lesions (n=5). **C-D**, As in **A**-**B** but for off-target lesions (n=7). **E**, Quantification of breathing and cries following bilateral on-target (Bilat.), unilateral on-target (Unil., n=6), and off-target (Off) electrolytic lesions. Each parameter is represented as a ratio of post-lesion:pre-lesion. Parameters are (from left to right) the total number of unisyllabic and multisyllabic (MS) cries (p = 0.002), proportion of cries that are MS (p = 0.006), the number of large breaths in a cry bout with a vocalization (p = 0.08), the length of each cry syllable (p = 0.01, 0.004, 0.002), and the basal respiratory rate (p = 0.1). Data are mean ± SEM. One-sided t-test or Wilcoxon rank sum P-value < 0.05 (*) and < 0.01 (**).

Neonates with bilateral on-target lesions still attempted to vocalize (ratio of post-lesion:pre-lesion cry bouts 0.86 ± 0.14 for on-target versus 1.08 ± 0.19 for off-target lesions, p = 0.2, t-test) by augmenting inspiratory and expiratory airflows (expiratory airflow: 0.89 ± 0.27 vs. 1.10 ± 0.07, p = 0.25 and inspiratory airflow: 0.92 ± 0.30 vs. 1.13 ± 0.13, p = 0.27), but failed to produce normal cries during these bouts and commonly exhibited irregular airflow (compare Fig. 4A with 4C). Bilateral on-target lesions resulted in a reduction in the number of unisyllabic and multisyllabic cries compared to off-target lesions and the few apparently normal augmented cry breaths were less likely to result in vocalizations (Fig. 4E). Furthermore, the very scarce successful cries were abnormally short following on-target versus off-target lesions (Fig. 4E). For each of these parameters, unilateral on-target lesions resulted in intermediate phenotypes (Fig. 4E). Importantly, basal breathing was unchanged in all lesioned animals (Fig. 4E). Moreover, impaired crying in neonates with on-target lesions could not be attributed to the ablation of TA or CT motor neurons, which reside more caudal and ventral to the area of lesion (*18*, *19*) (Fig. 3G, Fig. S3 and S4). Indeed, even off-target lesions that partially included the cNA resulted in normal crying. Together, these data demonstrate that the patterning of neonatal cries within a breath requires the rv-iRF, which contains premotor neurons for multiple vocalization articulators (Fig. 3), but that normal breathing is independent of this oscillator.

### A novel oscillator in the rv-iRF is active during expiration

We hypothesized that if the premotor cluster in the rv-iRF is the oscillator that controls fast rhythmic cries nested within a slower breathing pattern, its constituent neurons would oscillate faster than the preBötC inspiratory rhythm generator (*20*). We tested this idea using simultaneous recordings of ΔG-rabies-mCherry-positive TA premotor neurons in the rv-iRF and preBötC inspiratory activity in the hypoglossal nerve (cranial nerve XII) in medullary brainstem slices from neonatal mice (Fig. 5A). Whole cell recordings of premotor neurons revealed a rhythmic oscillation of membrane potential throughout expiration (occurring every 6.2 ± 1.6 s), which was faster than the rhythm of preBötC bursts (separated by 23.1 ± 1.8 s) (Fig. 5A and 5E). Furthermore, paired patch clamp recordings showed that individual neurons – either within a single rv-iRF or bilateral rv-iRFs – exhibited synchronous rhythmicity (Fig. 5B-C). To determine if this synchronous activity extends beyond the rv-iRF, we monitored the electrophysiological activity of a premotor neuron while simultaneously imaging GCaMP6s fluorescence emitted from all neurons in slices from *Snap25*-GCaMP6s transgenic mice (pan-neural expression of GCaMP6s). All neurons that displayed rhythmic changes in fluorescence during expiration were synchronized with each other and located in an area medial to the cNA. We found no evidence of rhythmic changes in GCaMP6s fluorescence in neighboring regions (Fig. 5D). This region precisely coincided with the rv-iRF containing the vocalization articulator premotor neurons, lesions of which nearly eliminated cries (Figs. 3 and 4). We named this cluster of dozens of neurons the intermediate reticular oscillator (iRO, Fig. 5D).

**Fig. 5.**
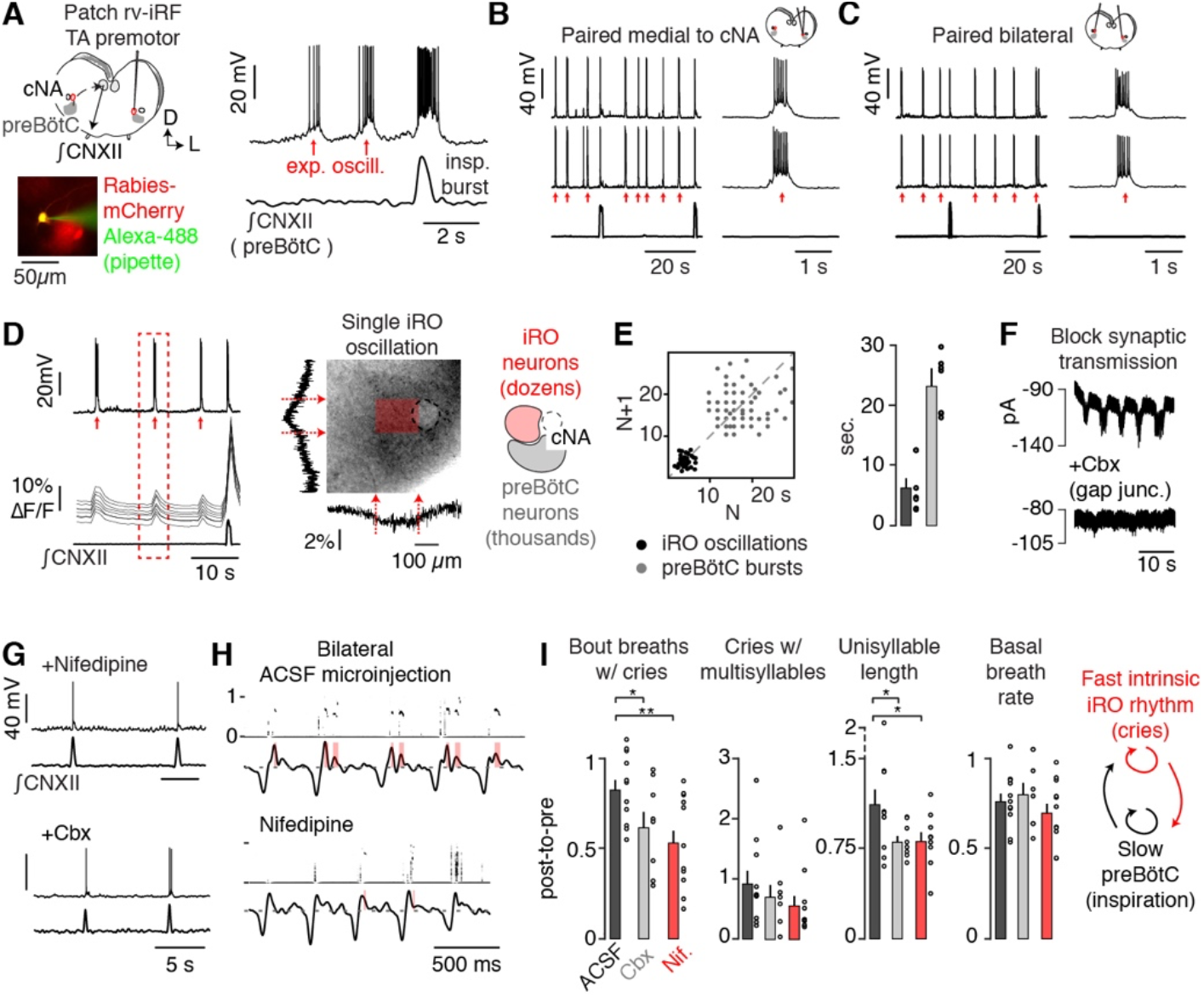
A novel oscillator in the rv-iRF is active during expiration. **A**, Left, medullary *in vitro* slice preparation. preBötC inspiratory rhythm monitored via cranial nerve XII (∫CNXII) and ΔG-rabies-mCherry TA premotor neurons in rv-iRF recorded using patch clamp (bottom, filled with Alexa-488 from pipette). Right, current clamp (I_c_) recording of TA premotor neuron showing rhythmical expiratory bursts between ∫CNXII activity, n = 17/20. **B**, Paired I_c_ recordings of rv-iRF neurons medial to cNA. Neurons show a synchronous expiratory oscillation. **C**, As in **B** but bilateral. **D**, Left, I_c_ recording of TA premotor rv-iRF neuron, ΔF/F in 10 neurons (*Snap25*-GCaMP6s), and ∫CNXII. Right, ΔF/F summed vertically and horizontally during a single expiratory oscillation (dashed box, left). ΔF/F localizes to dozens of neurons medial to cNA, denoted by shaded red box, which we named the iRO. **E**, Left, Poincaré plot of iRO (black) and preBötC (gray) intervals. Right, mean ± SEM (n=6). **F**, Voltage clamp recording of iRO neuron at −80 mV with synaptic blockers (10 μM NBQX, 50 μM APV, 100 μM Picrotoxin, 1 μM strychnine) then 100 μM carbenoxolone (n = 12). **G**, I_c_ recordings of iRO neurons with 2 μM nifedipine or 50μM carbenoxolone. The expiratory rhythm is abolished in both cases. **H**, Cry breaths after bilateral iRO injection of ACSF or 10μM nifedipine. **I**, As in Fig. 4E. Post-injection:pre-injection values for three vocalization parameters and basal breathing after microinjection of ACSF (n=11), nifedipine (n=11), or carbenoxolone (1mM, n=8). Data are mean ± SEM. One-sided t-test or Wilcoxon rank sum P-value < 0.05 (*) and < 0.01 (**).

Moreover, when fast synaptic communication between neurons was prevented with a cocktail of synaptic transmission blockers (10 μM NBQX, 50 μM APV, 100 μM picrotoxin, and 1 μM strychnine), all iRO neurons continued to oscillate synchronously (fig S7A, B). This uniquely defining feature further distinguished them from other nearby neurons. This iRO oscillatory activity was found in ΔG-rabies-traced TA (Fig. S5), CT, and tongue premotor neurons (Fig. S6A), but not premotor neurons for other orofacial oscillators (Fig. S6B) or respiratory centers in the brainstem (table S1). For example, although PiCo is nearby or perhaps overlapping, iRO is molecularly and electrophysiologically distinct (ChAT negative and does not require glutamatergic transmission) (*21*). We thus concluded that the iRO is a novel oscillatory node within the medullary brainstem, and it is uniquely composed of premotor neurons for multiple muscles of vocalization.

To investigate the mechanism underlying the oscillatory behavior of iRO neurons, we dissected the underlying ionic currents. In the presence of our cocktail of synaptic transmission blockers, all neurons retained synchronous rhythmic oscillations in membrane potential (Fig. S7A, B). Each iRO neuron displayed a rhythmically-timed inward current when the membrane potential was clamped at −80 mV, which was eliminated by the gap junction antagonists carbenoxolone (Fig. 5F), 18ß-glycyrrhetinic acid (Fig. S7C), and meclofenamic acid (Fig. S7C). Furthermore, the rhythmic activity in an iRO neuron persisted when voltage-gated sodium channels were blocked only within the patched neuron but was abolished when these sodium channels were blocked in nearby neurons (Fig. S7D). Importantly, the synchronized iRO neurons also showed direct electrical coupling (Fig. S7E). These data affirm that iRO neurons are coupled by gap junctions and that this, rather than fast synaptic signaling, is required for the autonomous oscillatory behavior that is unique to the iRO.

Using depolarizing and hyperpolarizing neuromodulators, we subsequently revealed that the iRO rhythm ceased upon network hyperpolarization and was therefore voltage dependent (Fig. S8A-B). This conditional rhythmic activity is consistent with the intermittent nature of crying. By analogy with other voltage-dependent rhythms, we predicted that at least three different currents would give rise to the interburst interval, the burst, and its termination. The canonical I_h_ pacemaker current carried by hyperpolarization-activated cyclic nucleotide-gated channels (*22*) was not involved in setting the interburst interval (Fig. S9A). However, block of the depolarizing persistent sodium current I_NaP_ eliminated the rhythm (Fig. S9A), consistent with a role in setting the interburst interval, burst, or both (*23*, *24*). Bursting was attenuated or prolonged by blocking or augmenting L-type calcium channel (LTCC) activity, respectively (Fig. S8C-D), but was not affected by a T-type calcium channel antagonist (Fig. S9A). Surprisingly, burst termination was not affected by any of the calcium-activated potassium channel antagonists that we tested (Fig. S9B), but was altered by preventing voltage-gated sodium channel inactivation (Fig. S8E).

This unique pharmacological profile (Fig. 5G) provided an opportunity to antagonize the iRO, but not the preBötC, rhythm *in vivo* to determine its requirement for patterning cries. To do so, we measured breathing and vocalizations before bilaterally injecting the iRO with either an LTCC or gap junction antagonist (nifedipine or carbenoxolone (Cbx), respectively), then remeasured breathing and vocalizations. Injection of Cbx or nifedipine significantly decreased the number of breaths with vocalizations in a cry bout and unisyllabic cry length, as well as reduced the proportion of cries with multiple syllables, (Fig. 5H-I, post-injection:pre-injection). In contrast, injection of ACSF as a control only minimally decreased the number of breaths containing vocalizations within a cry bout, and did not change the proportion of multisyllabic cries nor the length of unisyllabic cries (Fig. 5H-I). Consistent with our *in vitro* studies (Fig. 5G), basal breathing was unchanged by Cbx or nifedipine (Fig. 5I). The changes observed in cries after antagonizing the iRO rhythm mirrored those produced by electrolytic lesions, and thus demonstrated that the iRO’s intrinsic rhythm is required for patterning neonatal cries, consistent with it being the cry oscillator (Figs. 1 and 2).

### The iRO recruits the preBötC and laryngeal motor neurons to produce the cry motor program

The first feature of the cry motor program is the activation of inspiratory muscles that precedes the onset of each cry syllable (Figs. 6A and 2). We postulated that the iRO triggers the inspiratory motor program via the preBötC. *In vitro*, ~50% of recorded preBötC neurons displayed excitatory postsynaptic potentials (EPSPs) during each iRO oscillation (Fig. 6B), demonstrating excitatory synaptic connections from the iRO to the preBötC. Furthermore, preBötC bursts occurred in phase with, or immediately after, the onset of iRO oscillations (Fig. 6C and Fig. 6D, compare 2 vs. 3), showing that the iRO dictates the timing of the preBötC bursts. Reciprocal excitatory connections from the preBötC reset the iRO rhythm after each inspiration (Fig. 6D, compare 1 vs. 2). Thus, the iRO has the connectivity necessary to initiate the inspiratory motor program that precedes each of the cry syllables within a breath (Fig. 6D).

**Fig. 6.**
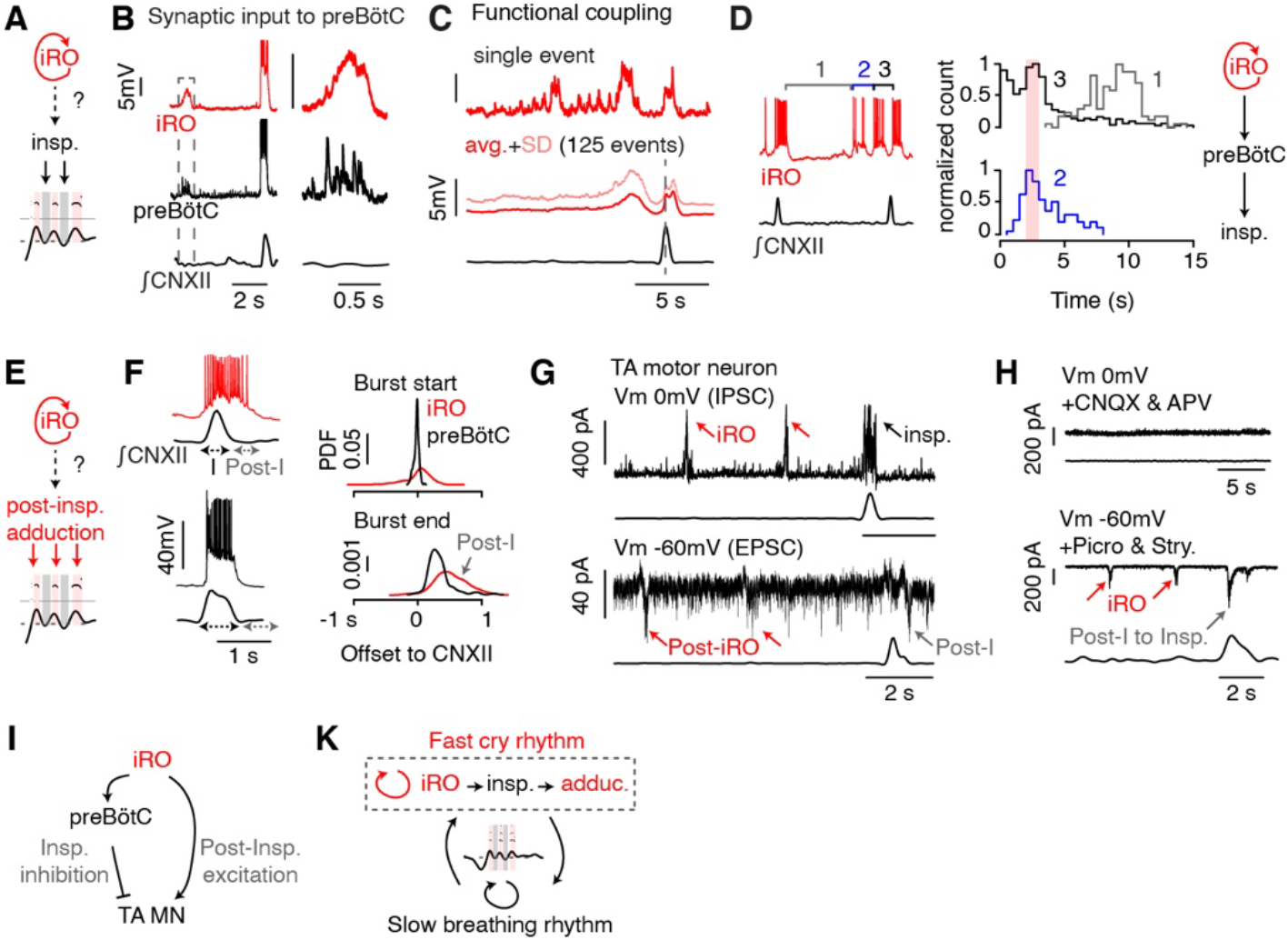
The iRO uses synaptic and functional connections to produce the cry motor program. **A**, Schematic of iRO control of inspiration (Fig. 2). **B**, Left, paired I_c_ recordings of iRO and preBötC inspiratory neurons (n = 5/11 paired recordings). PreBötC burst is concurrent with ∫CNXII activity and iRO oscillation is without. Right, magnified boxed portion showing EPSPs in preBötC inspiratory neurons during iRO expiratory oscillation. **C**, Top, 15 second iRO I_c_ recording and bottom, average membrane potential ± SD for 125 events aligned to ∫CNXII peak. Action potentials were removed to better visualize membrane potential oscillations. **D**, Left, schematic showing segments measured: 1 - preBötC burst to first iRO oscillation (gray), 2 - intervals between expiratory iRO oscillations within a cry breath (blue), 3-last iRO oscillation to preBötC burst (black). Right, histogram of duration of each segment (1: n = 134 events, 2: n = 182 events, 3: n=715 events). Data from 13 mice. **E**, Schematic of iRO induction of post-inspiratory laryngeal adduction (Fig. 2). **F**, Left, examples of iRO (red) and preBötC (black) bursts with associated ∫CNXII activity. iRO activity persists into post-inspiration (Post-I). Right, probability density function showing distribution of burst start and end compared to peak ∫CNXII activity (n = 7 iRO and 12 preBötC neurons). **G**, TA motor neurons identified by intramuscular injection of cholera toxin B-555. Top, example inhibitory post-synaptic currents (IPSCs) in 22/23 motor neurons held at a membrane potential (V_m_) of 0 mV. Bottom, example excitatory post-synaptic currents (EPSCs) in 13/22 motor neurons held at −60 mV. Both input types coincide with iRO oscillations and EPSCs are post-inspiratory. **H**, Top, IPSCs are absent in CNQX and APV (V_m_, 0 mV). Bottom, post-inspiratory EPSCs are now associated with inspiration in picrotoxin and strychnine (V_m_, −60 mV), mirroring iRO activity. **I**, Schematic of TA inputs that pattern post-inspiratory activity. **K**, Summary model depicting rhythmic induction of a motor program by the iRO. Syllables are produced by a fast rhythm nested within slower breathing.

The second key feature of the cry motor program is the laryngeal adduction to produce sound that follows each inspiratory event, known as post-inspiratory activity (*25*) (Fig. 6E). In contrast to preBötC neurons, the excitation of iRO neurons during inspiration persisted through post-inspiration (Fig. 6F). Because iRO neurons are glutamatergic and premotor to laryngeal adductors, their post-inspiratory activity is consistent with the ability to generate cries. However, an additional mechanism must be invoked to explain the prevention of laryngeal adduction during inspiration. We investigated this using voltage-clamp recordings of TA motor neurons *in vitro* following their identification by intramuscular injection of Alexa Fluor 555-conjugated cholera toxin B (Fig. 6G-H). Nearly all recorded TA motor neurons exhibited inhibitory synaptic currents during each inspiration, and 60% displayed excitatory post-inspiratory activity, which we expected to result from the iRO activity (Fig. 6G). These excitatory and inhibitory inputs were also evident during each iRO oscillation (n = 12/22 motor neurons) (Fig. 6G). This *in vitro* activity mirrors the timing of laryngeal sound production during each cry syllable.

Blockade of excitatory synaptic transmission (using 10 μM CNQX and 50 μM APV) silenced the preBötC (*20*) and even eliminated all inhibitory synaptic modulation of TA motor neurons (Fig. 6H). As iRO remains rhythmic in this condition (Fig. 5F and Fig. S7A), the direct inhibitory input to TA motor neurons must therefore be from a different source, likely the Vgat-positive TA premotor neurons within the preBötC or RAm (Fig. 3). Upon blockade of inhibitory synaptic transmission (using 100 μM picrotoxin and 1 μM strychnine), excitatory synaptic input remained, and accurately mimicked iRO activity, as expected (Fig. 6H vs. 6F). This connectivity between the iRO, the inhibitory inspiratory neurons, and TA motor neurons comprises a plausible mechanism by which iRO patterns post-inspiratory laryngeal adduction and sound production (Fig. 6I). Thus, by recruiting the preBötC to drive the inspiratory motor program and directly activating laryngeal adductors post-inspiration, the iRO patterns neonatal cries.

## Discussion

We have identified a novel brainstem node, the iRO, which acts as an autonomous oscillator that organizes the patterning of neonatal mouse vocalizations with breathing. Its proximity to the cNA is consistent with a previous hypothesis that a central pattern generator for vocalization resides in the medullary reticular formation (*6*). A simple model predicts that patterning of laryngeal and breathing motor groups is sufficient to control vocalizations (*7*). Furthermore, it is anticipated that such a patterning system would also encode the tempo of syllables within vocalizations. Our results are consistent with such a model.

We found that neonatal cries contain one or more regularly spaced syllables. The production of sound requires adduction of the larynx by the TA and each syllable is separated by reactivation of inspiratory muscles. Thus, the standard vocalization motor program – inspiration followed by laryngeal adduction – must be repeatedly engaged within a single breath to generate multisyllabic cries. Our identification and characterization of the iRO, which produces a fast and autonomous rhythm nested within a slower breathing rhythm, provides a mechanism for such repetition. Each burst of iRO activity patterns the motor program to enable articulation of a single cry, and multiple iRO bursts within a breath generate multisyllabic cries (Fig. 6K). Ablation or pharmacological inhibition of the iRO either eliminates or modifies normal neonatal cry rhythms and disrupts multisyllabic cry production. Any residual, albeit abnormal, cries and airflow augmentation are likely generated by another brainstem region, such as the RAm (*26*). It will be important to determine whether the iRO’s role in timing syllable onset is generalizable to other mouse vocalizations, as well as those of different species, and how this is utilized or bypassed during learned vocalizations or song.

Hardwired patterning systems like the iRO allow the brain to generate robust, reproducible motor tasks that are essential for innate behaviors, such as crying. Because the iRO can coordinate breathing with laryngeal closure, we imagine that it may be repurposed for non-vocalization behaviors that require laryngeal closure, including spontaneous laughter (*27*, *28*) and swallowing (*29*–*31*). Indeed, neurons within the anatomical region of the iRO have previously been implicated in swallowing (*29*). Both its restricted anatomical location and rhythmic activity make the iRO an ideal system for studying the role of higher brain centers and neuromodulation in sculpting or repurposing a pattern generating system to support a range of both innate and learned mammalian behaviors.

## Acknowledgments

We thank Dr. Eric Lam (University of California, San Francisco Kavli-PBBR Fabrication and Design Center) for his assistance in design and fabrication of the plethysmograph. We thank Adelae Durand (University of California, San Francisco) for her assistance in neonatal plethysmography recordings. We thank Dr. David Julius (University of California, San Francisco), Dr. Massimo Scanziani (University of California, San Francisco), Dr. Roger Nicoll (University of California, San Francisco), and members of the Yackle lab for their input and revision of the manuscript.

## Author Contributions

X.P.W. designed and developed the neonatal plethysmograph and ultrasonic recording system. X.P.W. and K.Y. collected and analyzed vocalization, breathing, and EMG data. X.P.W. performed laryngeal microinjections, stereotaxic brain injections, and electrolytic lesions. X.P.W. and B.D. performed rabies tracing and immunohistochemical studies. X.P.W., M.C., and K.Y. performed and analyzed electrophysiological and calcium imaging studies. B.D. and G.F. originally designed independently conducted laryngeal rabies virus tracing studies. X.P.W. and K.Y. wrote the manuscript and all authors edited the manuscript.

## Data and materials availability

All data collected in this study and code use for analysis are available upon request from the corresponding author.

