## Supplementary Materials for "A novel reticular oscillator in the brainstem synchronizes neonatal crying with breathing"

### Materials and Methods

Animals

C57Bl/6J, *Snap25-GCaMP6s* (*Snap25*$^{tm3.1Hze}$) (*32*), *Slc17a6*-Cre (*33*), *Gad2*-Cre (*34*), *Slc32a1*-Cre (*33*), *Slc6a5*-Cre (*35*), *RIKEN-Slc6a5*-Cre (*36*), *Dbx1*-Cre (*37*), *Foxp2*-Cre (*38*), *Chat*-Cre (*39*), *Egr2*-Cre (*40*), *Neurod6*-Cre (*41*), *Penk*-Cre (*42*), *Parv*-Cre (*43*), *Tac1*-Cre (*44*), *Npy*-Cre (*45*), ROSA-LSL-G-TVA (Gt(ROSA)26Sor$^{tm1(CAG\text{-}RABVgp4,\text{-}TVA)Arenk}$) (*46*), ROSA-LSL-GCaMP6s (Ai96) (*32*), ROSA-LSL-GCaMP6f (Ai95) (*32*), ROSA-LSL-EYFP (*47*) have been described. Mice were housed in a 12-hour light/dark cycle with unrestricted food and water. All animal experiments were performed in accordance with national and institutional guidelines with standard precautions to minimize animal stress and the number of animals used in each experiment.

Recombinant viruses

All viral procedures followed the Biosafety Guidelines approved by the University of California, San Francisco (UCSF) Institutional Animal Care and Use Program (IACUC) and Institutional Biosafety Committee (IBC). The following viruses were used: oG SiR G-Deleted Rabies-FlpO-mCherry, G-Deleted Rabies-eGFP, G-Deleted Rabies-mCherry (all >1.0x$10^8$ TU/mL, The Viral Vector Core at the Salk Institute for Biological Sciences or Janelia Viral Tools) and HSV-hEF1a-Rabies G (RN700), HSV-hCMV-YTB (Gene Delivery Technology Core Massachusetts General Hospital).

Plethysmography, vocalization, and analysis

Breathing pressure changes and vocalizations were measured in neonatal mice (P2-P4) in a custom prepared whole animal plethysmography chamber at room temperature (23°C, Figure 1a) and constant humidity. The top of the chamber is sealed by the microphone and enables the volume of the chamber to be appropriately adjusted to the size of the neonate. Pups were removed from their home cage and placed in the recording chamber for 2 x 5 minutes of recording and returned to their home cage. Mice were recorded for 1-2 more times throughout day, following the same protocol, to get a consistent baseline. Vocalization was recorded using an Avisoft Bioacoustics UltraSoundGate CM16/CMPA microphone and chamber pressure was transmitted to a spirometer (AD Instruments). The microphone and spirometer were connected to a PCI DAQ device (PCI NIDAQ 6251), allowing respiration and vocalization data to be simultaneously acquired at 400kHz using custom Matlab (MathWorks) code. Light tapping of the chamber with a metal rod show a constant <2ms difference in timing of changes in sound and pressure change.

The respiration and vocalization data were analyzed offline in Matlab. The pressure trace was downsampled, filtered, and breathing was segmented using the troughs and peaks of measured pressure as surrogates for onset of inspiration and expiration, along with quality control metrics. Pressure change was defined as the first derivative of the pressure trace with respect to time. Given differences in tidal volume due to the intrinsic variance of lung volumes of pups at these ages (discussed further below), large breaths were defined as the minimal tidal volume that must be exceeded for 95% of cries in a trace. USVs were detected using code modified from the Holy Lab (*48*). The parameters used to detect vocalizations are: spectral purity threshold > 0.3, spectral discontinuity threshold < 0.8., cry duration threshold > 2ms, minimal inter-syllable interval > 15ms, and mean frequency threshold > 30kHz.

Since barometric whole body plethysmography imprecisely measures airflow and tidal volume (*49*), tidal volume in neonates correlates with their weight instead of age (*50*), we have reported the breathing measurements as changes in pressure that are in arbitrary units. Thus, we have not compared the amplitude of breathing airflow between animals nor reported it as mLs/sec. However, these measurements still allow for the changes in breathing within the same animal to be directly compared as arbitrary units.

Genioglossus and intercostal EMG

Neonatal mice were briefly anaesthetized by hypothermia. Paired EMG electrodes made from Teflon-coated multistranded stainless steel wires (Cooner Wire) and small suture needles (Fine Science Tools) were passed through the genioglossus or intercostal muscles and secured with a knot (*51*). Teflon was removed from a small (<0.5mm) segment of the wire adjacent to the knot, allowing the exposed wire to be embedded in the GG or intercostal muscles. The EMG electrodes were connected to an AC differential amplifier (A-M systems AM1800), band-pass filtered (300Hz/10kHz) and digitized with an PCI DAQ device (PCI NIDAQ 6251), along with USV and respiration measurements, allowing for synchronous data acquisition at 250kHz (single EMG) or 200kHz (dual EMGs). The EMG data was rectified and integrated offline with a modified Paynter filter in Matlab.

Laryngeal (TA) Muscle Lesion

Neonatal mice were anaesthetized by hypothermia and gently fixed in the supine position. A small 2mm incision was made ventrally in the neck, and the muscles overlying the thyroid cartilage were carefully isolated with sutures. A microincision was made on the thyroid cartilage with the sharp tip of a 28G needle, and a small bipolar electrode (Microprobes) was inserted into the TA muscle. The muscle was lesioned by passing a current of 500μA for 2.5-5 seconds. For the sham group, the same protocol was followed except no current was passed after inserting the bipolar electrodes into the TA muscle. The neck was sutured, and mouse was allowed to recover fully before recording of attempted vocalizations. After the recording, the larynx was fully dissected to confirm the site of the lesion.

Stereotaxic electrolytic lesion and injection

Bilateral stereotaxic injections and electrolytic lesions were performed in neonatal mice anaesthetized by hypothermia. The mouse was oriented on a stereotaxic frame (Kopf) with a neonatal adaptor (Stoelting) such that bregma was 1.5mm below lambda. In this position, the coordinates used for the iRO node were: P2: 2.50 mm posterior, 3.35 mm ventral from surface of the brain, ± 0.82 mm lateral from lambda; P3: 2.30 mm posterior, 3.50 mm ventral from surface of the brain, ±0.87 mm lateral from lambda. Electrolytic lesions were performed with a concentric bipolar microelectrode (FHC) by passing a current of 30-80μA for 2-5 seconds. Animals were fully recovered on a heat pad and returned to their nest. After lesions, neonates recovered for at least 24 hours before breathing and vocalizations were recorded again. After the post lesion recordings were performed, lesion sites were validated by both anatomical localization visualized by light microscopy and changes of GCaMP6s activity in the iRO and preBötC in a medullary slice preparation (described below). Lesions were classified as bilateral “off” target from the iRO node, bilateral “on” target, or unilateral “on” target. “off” target lesions were used as the control group and therefore all breathing, vocalization, and physiology validation were performed as a blinded study. The same coordinates were used for stereotaxic injection. Approximately 70nL of nifedipine (10μM), 70nL of carbenoxolone (1mM), or 70nL of ACSF were injected with a Nanoject III (Drummond), at a rate of 2-3nLs/second. The animals were recovered on a heating pad and breathing and vocalization were recorded ~30 minutes after injection. Injection sites were confirmed by histological analysis of co-injected dye.

Laryngeal Rabies virus and Cholera toxin injections

Rabies tracing was performed using Chat-Cre;ROSA-LSL-G-TVA (Gt(ROSA)26Sor$^{tm1(CAG-RABVgp4,-TVA)Arenk}$) mice and the G-Deleted Rabies viruses listed above. For a subset of tracing experiments, a G-Deleted Rabies virus was coinjected with HSV hEF1a-Rabies G in C57Bl/6J or *Snap25*-Gcamp6s mice. Neonatal mice at P0 were cryoanesthetized and fixed in various positions with adhesive that were suitable for the muscle being injected. Each muscle was injected using a Nanoject III (Drummond). Between 30 to 150 nLs (depending on muscle) of virus were injected unilaterally, depending on the muscle, at a rate of 5-10nL/sec. Muscles injected were the cricothyroid, thyroarytenoid, genioglossus, masseter, whisker pad, and nose in order to trace the premotor neurons for vocalization (larynx, genioglossus), swallowing (genioglossus), chewing (masseter), and whisking (whiskers), and nose movement (nose). For injection into the laryngeal muscles (thyroarytenoid and cricothyroid), the overlaying muscles were bluntly dissected and carefully isolated with sutures. A microincision was made on the thyroid cartilage with the sharp tip of a 28G needle to allow for insertion of the micropipette into the thyroarytenoid muscle. Normal saline was used to flush the overlying area to prevent any nonspecific infection and labeling. After injections, the neonates were recovered on a heat pad and then returned to their nest. Neonates recovered for 4 days and then were euthanized for medullary slice preparation (described below). Rabies traced fluorescent neurons were identified and their electrophysiological activity was recorded. Cholera Toxin Subunit B (C34776, Life Technologies) injections were performed as described above, except they were only into the thyroarytenoid and cricothyroid muscles, and animals recovered for 2 days before euthanasia for brainstem slice preparation. All injection solutions contained fast-green dye to confirm that only the correct muscle was injected.

Histology

4-7 days after injection of the viral cocktail, pups were transcardially perfused with 10mL of heparinized saline followed by 10mL of 4% PFA in phosphate-buffered saline (PBS). The brains were dissected and postfixed overnight in 4% PFA, cryoprotected in 15 or 30% sucrose PBS (w/v) and embedded in OCT. The embedded brains were cryosectioned at 20 or 30µm. Sections were then blocked for 1 hour in 0.5% triton-X PBS and 4% normal horse or goat serum and subsequently incubated with a solution of primary antibodies in the blocking solution at 4°C overnight. For samples that required the use of a mouse primary antibody, the sections were further blocked using the Mouse on Mouse Blocking Reagent (Vector Labs, MKB-2213-1) for 1 hour prior to incubation in the primary antibody. The primary antibodies used are: chicken anti-bGal (Abcam, ab9361), chicken anti-GFP (Aves, GFP-1020), goat anti-ChAT (Millipore, AB144p), rabbit anti-GFP (Invitrogen, A11122), rabbit anti-phox2b (Jean-François Brunet lab), mouse anti-phox2b (Santa Cruz, B-11), rat anti-RFP (Chromotek, 5F8). The sections were washed in PBS, and the incubation procedure was then repeated with a solution containing secondary antibodies for 1-4 hours at 4°C. The secondary antibodies used are: donkey anti-chicken 488 (Jackson laboratories, 703-545-155), donkey anti-chicken Cy5 (Jackson laboratories, 703-176-155), donkey anti-goat Cy5 (Jackson laboratories, 705-606-147), donkey anti-rabbit 488 (Jackson laboratories, 711-545-152) donkey anti-rabbit Cy5 (Jackson laboratories, 712-165-153), donkey anti-rat Cy3 (Jackson laboratories, 711-495-152), goat anti-mouse Alexa Fluor 647 (Invitrogen, 21236). The slides were washed in PBS, air dried and mounted using mowiol or fluorescence mounting medium (Dako) and coverslipped. Epifluorescence and confocal images were acquired with a NanoZoomer S210 digital slide scanner (Hamamatsu Photonics), Leica SP5 confocal microscope (Leica), or Nikon CSU-W1 Spinning Disk (Nikon).

Slice preparation and electrophysiology

550 to 650μm-thick transverse medullary slices which contain the preBötC and cranial nerve XII (XIIn) were prepared from neonatal P0-5 *Snap25*-GCaMP6s, *Chat-Cre;*ROSA-LSL-G-TVA (Gt(ROSA)26Sor$^{tm1(CAG-RABVgp4,-TVA)Arenk}$) rabies injected, HSV and rabies injected, and lesioned animals were prepared as described (*52*). Briefly, slices were cut in ACSF containing (in mM): 124 NaCl, 3 KCl, 1.5 $CaCl_2$, 1 $MgSO_4$, 25 $NaHCO_3$, 0.5 $NaH_2PO_4$, and 30 D-glucose, equilibrated with 95% $O_2$ and 5% $CO_2$ (4°C, pH=7.4). The rostral side of the slice was taken 100μm caudal to the end of the facial nucleus, at the rostral end of the compact nucleus ambiguus. All recordings were performed in the ACSF described above except the $K^+$ was raised to 9 mM and temperature to 27.5-28.5°C. This elevation of extracellular $K^+$ enables spontaneous preBötC activity to activate the hypoglossal motor nucleus (*53*). The preBötC neural activity was inferred by the activity recorded from either XIIn rootlet or as population activity directly from the XII motor nucleus using suction electrodes, amplified and low/high pass filtered at 3kHz/400Hz, rectified, integrated and digitized using Digidata 1550B. Current and voltage clamp recordings of single neurons were performed with a MultiClamp700A or B using pClamp9 and digitized at 10000 Hz. Internal consisted of $K^+$-Gluconate (135mM), EGTA (1.1mM), NaCl (5mM), $CaCl_2$ (0.1mM), HEPES (10mM), ATP (2mM), GTP (0.3mM). Alexa-488 was used to fill recorded neurons. For experiments in synaptic blockers, NBQX (10μM, Abcam, ab120046) or CNQX (10μM, Abcam, ab120044), D-APV (50μM, Alomone Labs, D-145), picrotoxin (100μM, Abcam, ab120315),

strychnine (1µM, Sigma Aldrich, S0532) were bath applied after an initial 20-minute recovery period in 9mM $K^+$ ACSF. For pharmacological analysis of rhythmic activity, the following peptides and drugs were bath applied: Carbenoxolone (50-150µM, Sigma-Aldrich), 18ß-glycyrrhetinic acid (150µM, Sigma-Aldrich, G10105), Meclofenamic acid (100µM. Fisher Scientific, AAJ6048403), tetrodotoxin (TTX, 1µM, Abcam, ab120054), Substance P (200nM, Abcam, ab120170), DAMGO (20-200nM, Abcam, ab120674), Nifedipine (2µM, Tocris, 1075), Bay K8644 (100nM, Tocris, 154410), Veratridine (250nM, Abcam, ab120279), ZD7288 (100µM, Abcam, ab120102), Riluzole (20µM, Abcam, ab120272), Mibefradil (10µM, Tocris, 219810), Iberiotoxin (50nM, Alomone Labs, STI-400), UCL1684 (50nM, Tocris, 13105). For pharmacological analysis of rhythmic activity, the drugs were added to the internal recording solution: QX314 chloride (5mM, Tocris, 2313).

Gap junction electrical coupling

Gap junctions between the iRO neurons were validated by both electrical coupling and dye coupling. To determine if neurons were electrically coupled, the iRO neurons were identified by Snap25-GCaMP6s activity in 9mM $K^+$ ACSF containing synaptic blockers. After a pair of iRO neurons were patched, TTX was bath applied to silence the spontaneous rhythm. A current step of 500pA was applied to one neuron and a corresponding change in membrane potential was recorded. After, a similar current protocol was applied to the alternative neuron.

Statistics

Statistical tests were performed on data collected from electrolytic lesions, nifedipine, ACSF, and carbenoxolone stereotaxic injection. For all datasets, a Shapiro-Wilks test was first performed to assess normality of the distribution. P-value <0.05. If both groups to be compared had normal distributions, a one tailed t-test with unequal variance was performed. If one of both of the data sets were not normal, then a Wilcox Rank Sum was performed. For electrolytic lesions, the t-test compared "off" target to bilateral "on" target lesions and “off” target to unilateral lesions. For pharmacology microinjections, ACSF was compared with nifedipine and carbenoxolone.

### Data Availability

All data collected in this study is available upon request from the corresponding author.

### Code Availability

All analysis code developed in this study is available upon request from the corresponding author.

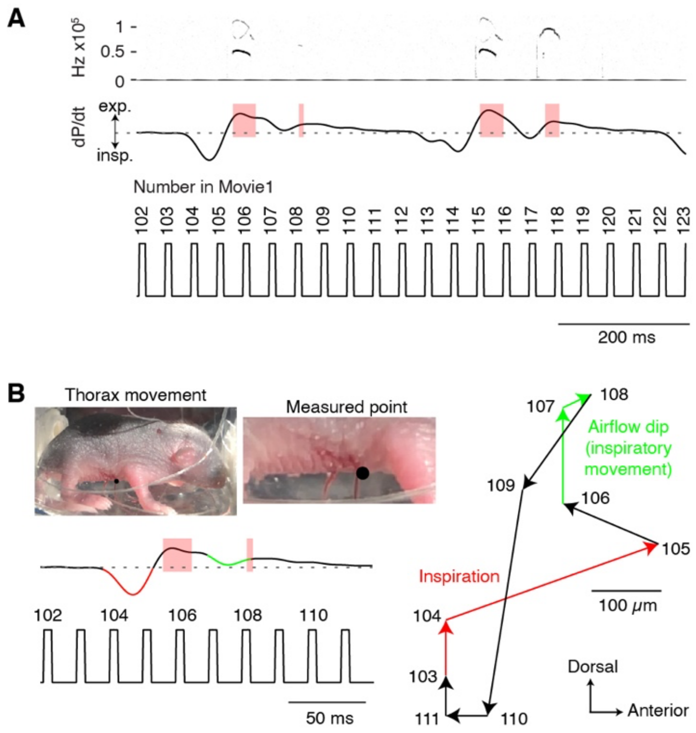


**Fig. S1. Breathing pressure changes and cries that correspond to the breathing movements in Movie S1.** **A**, Spectrogram and respiratory pressure changes for two sequential breaths with bisyllabic cries. Numbers below correspond to Movie S1 and are meant for alignment. Inspiration occurs in frames 103-105 and 113-115. Inspiratory-like events that separate each syllable occur in frames 106-107 and 116-118. **B**, Quantification of the thoracic body wall movements in Movie S1. Black dot on the thorax represents the point tracked during the breath. As inspiration occurs, 103-105 (red), the thorax retracted (103-104) and moved anterior (104-105). This same movement, although smaller in magnitude, occurred during the inspiratory like event between the two syllables 106-108 (green). During expiration, the body wall relaxed (108-111). These body wall movements are consistent with the inspiratory motor program measured in Fig. 2 between syllables and demonstrates that the airflow oscillation between two syllables is likely generated by an inspiratory motor program.

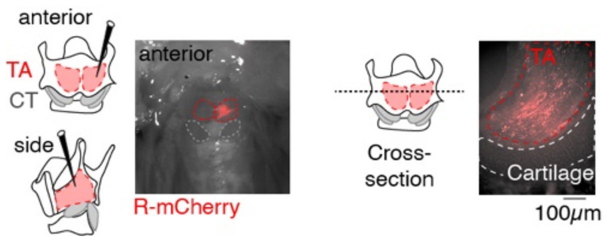


**Fig. S2. Confirmation that rabies virus injection is restricted to the thyroarytenoid muscle.** Visualization of the motor neuron axons within the thyroarytenoid muscle 4-7 days after unilateral (ΔG-rabies-mCherry microinjection. Left, anterior view of the larynx. Dashed red outline, thyroarytenoid muscle (TA). Dashed gray outline, cricothyroid muscle (CT). Note fluorescence is restricted to the single TA muscle that was injected. Right, cross section of the larynx showing motor neuron axons underneath the thyroid cartilage in the TA muscle.

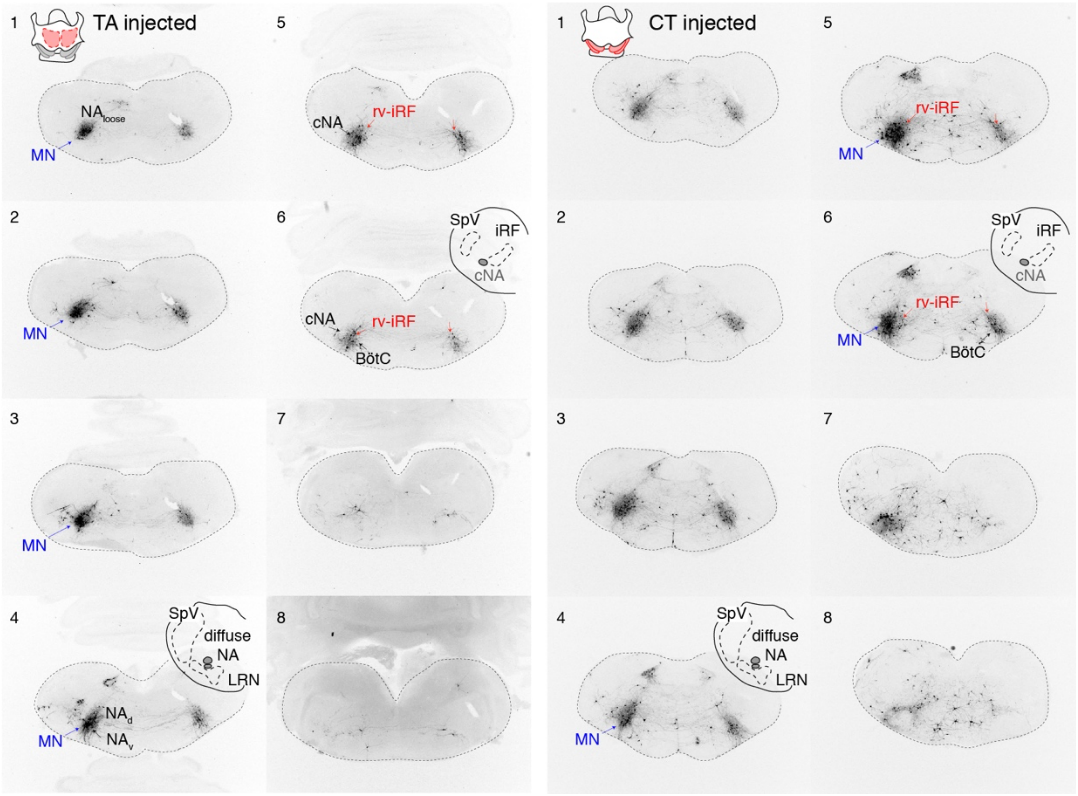


**Fig. S3. Complete medullary motor and premotor pattern from rabies-traced laryngeal muscles.** Sequential coronal sections of the medullary brainstem after ΔG rabies-tracing initiated unilaterally from the thyroarytenoid (TA, left) and the cricothyroid (CT, right) muscles. Sections are organized from caudal (1) to rostral (8). Red arrows in sections 5 and 6 indicate the anatomical location of rv-iRF. Note, rv-iRF neurons are premotor since they do not express Phox2b (Fig. 3). Blue arrows indicate traced motor neurons (MN). These are identified by their large size and Phox2b expression (Fig. 3, fig. S4). Furthermore, their locations match the MN patterns identified by HSV-GFP (fig. S4), the absence of Vglut2 and Vgat expression (Fig. 3), and the literature (*18*). Black arrows indicate the loose formation of the nucleus ambiguus ($NA_{loose}$), semicompact ($NA_d$ and $NA_v$), and compact (cNA). Black arrows on section 6 also indicate BötC. SpV, Spinal trigeminal nucleus. iRF, intermediate Reticular Formation.

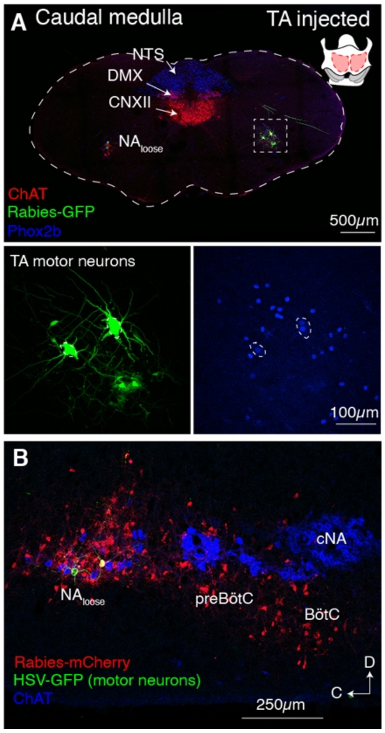


**Fig. S4. Anatomical location of thyroarytenoid motor neurons.** **A,** Example of coronal brainstem slice showing the thyroarytenoid (TA) motor neurons. Magnifications of boxed region below. Motor neurons are identified by colocalization of ΔG rabies-GFP (green) Choline acetyl transferase immunostain (ChAT, red), and Phox2b immunostain (blue, shown below). Note, the TA motor neurons are in the $NA_{loose}$ (consistent fig. S3) in the caudal brainstem where the NTS (nucleus tractus solitarius, Phox2b positive), DMX (dorsal motor nucleus 10, Phox2b and ChAT positive), and CNXII (cranial nerve 12, ChAT positive) are located. **B**, Sagittal section of the lateral medulla showing TA traced motor neurons identified by HSV-Glycoprotein-GFP and ChAT expression (green) and ΔG rabies-traced premotor neurons (red, rabies-mCherry positive, HSV-GFP negative, and ChAT negative). TA motor neurons are located in the $NA_{loose}$. C, caudal. D, dorsal.

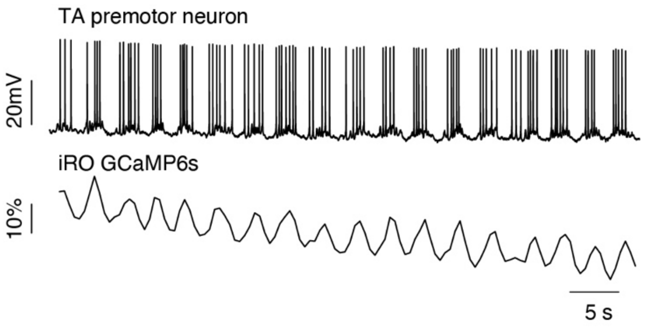


**Fig. S5. Thyroarytenoid premotor neurons are part of the iRO gap junction dependent rhythm.** Current clamp recordings of a ΔG rabies-traced thyroarytenoid premotor neuron in *Snap25*-GCaMP6s mice with bath applied blockers of fast synaptic neurotransmission (10μM NBQX, 50μM APV, 100μM Picrotoxin, 1μM Strychnine). The premotor neuron has rhythmic activity that is synchronous with the iRO oscillation in GCaMP6s fluorescence. ΔF/F scale bar, 10%.

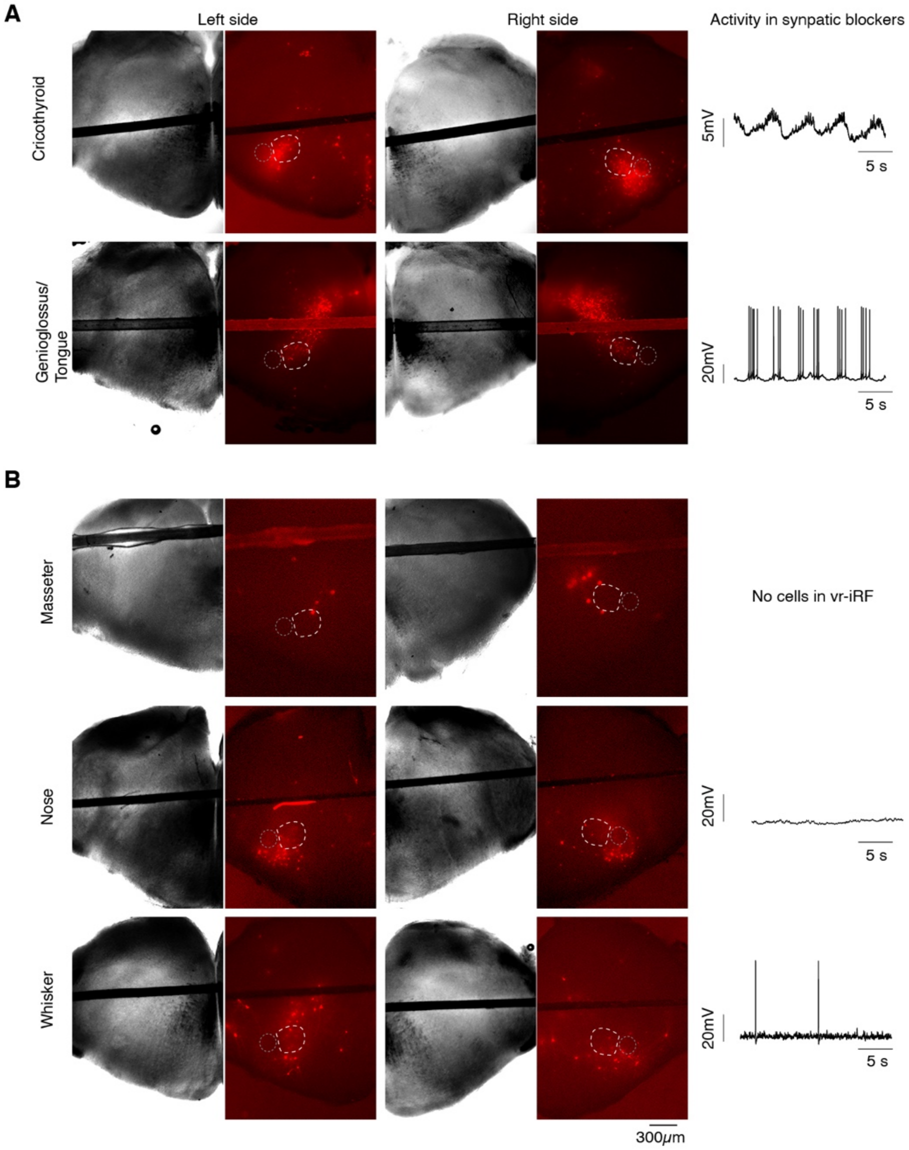


**Fig. S6. Anatomical location of rabies-traced premotor neurons from other orofacial muscles.** **A**, Left, Corresponding light and fluorescent microscopy images of the left and right sides of medullary brainstem slices depicting the location of the ΔG rabies-mCherry traced motor and premotor neurons (red) after injection of rabies into the cricothyroid and tongue. rv-iRF (dashed white), cNA (dashed gray). Right, Changes in membrane potential measured in current clamp of ΔG rabies neurons in the rv-iRF in fast synaptic neurotransmission (10μM NBQX, 50μM

APV, 100µM Picrotoxin, 1µM Strychnine). Oscillation in membrane potential indicates these neurons are within the iRO network (n = 7/19 for CT, n = 5/20 for tongue). **B**, As in **A**, but ΔG rabies-mCherry injected into other orofacial oscillators. Masseter (chewing, no neurons in rv-IRF), nose (sniffing, n = 0/6), and whisking (whisking, n = 0/5). The few neurons within rv-iRF did not display oscillations in membrane potential. Consistent with the coordination of nose movement and whisking with respiration, some rabies traced neurons localized to the preBötC/BötC, as has previously been described (*54–56*).

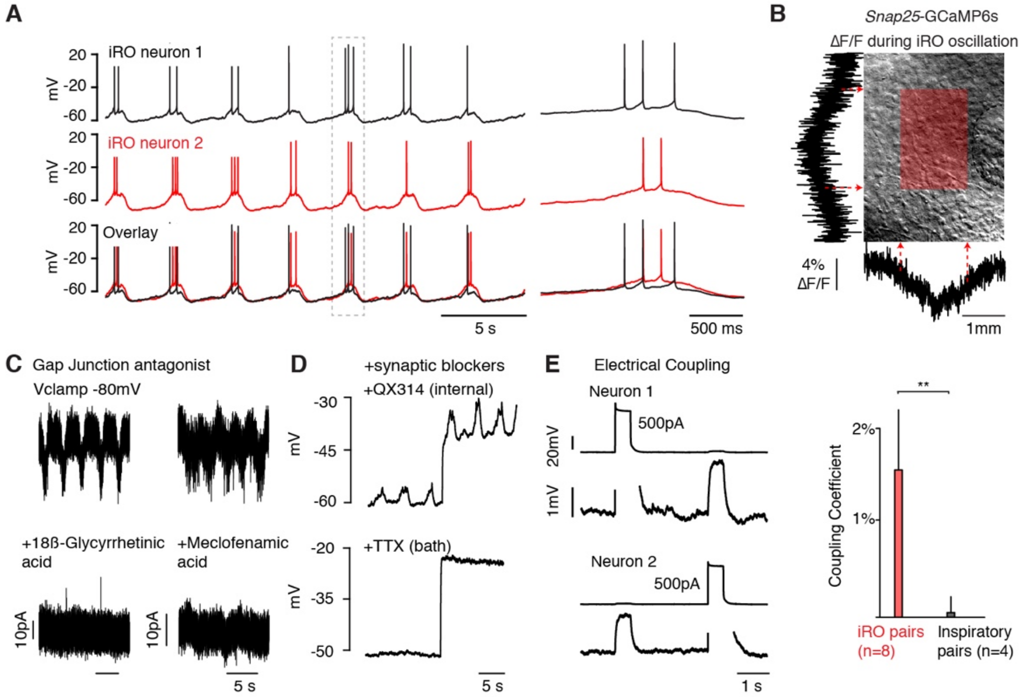


**Fig. S7. Multiple experiments confirm gap junctions between iRO neurons and the requirement for the iRO rhythm.** **A**, Paired current clamp recordings of two iRO neurons in blockers of all fast synaptic neurotransmission (10µM NBQX, 50µM APV, 100µM Picrotoxin, 1µM Strychnine). N = 16. **B**, ΔF/F summed vertically and horizontally during a single oscillation in synaptic blocker. ΔF/F localizes to only dozens of neurons medial to cNA. **C**, -80mV voltage clamp recordings before (top) after (bottom) bath application of gap junction antagonists 18ß-glycyrrhetinic acid (150µM, n = 2) and meclofenamic acid (100µM, n = 4). All recordings in fast synaptic blockers. **D**, Current clamp recording in fast synaptic blockers. Top, internal solution with QX314 (5mM). Bottom, same cell after bath application of tetrodotoxin (bottom, 1µM, n = 4). **E**, Paired current clamp recordings in synaptic blockers and 1µM TTX. Large current steps in one neuron could be measured as several mV changes in membrane potential in the second neuron. Right, quantification of coupling coefficient for control paired recordings and iRO neurons. N=5-

20 trial per paired recording. iRO pair coupling coefficient 1.2 ± 1.4%, n = 8; control non-iRO pairs nearby 0.00 ± 0.6%, n = 4. T-test P-value = 0.007.

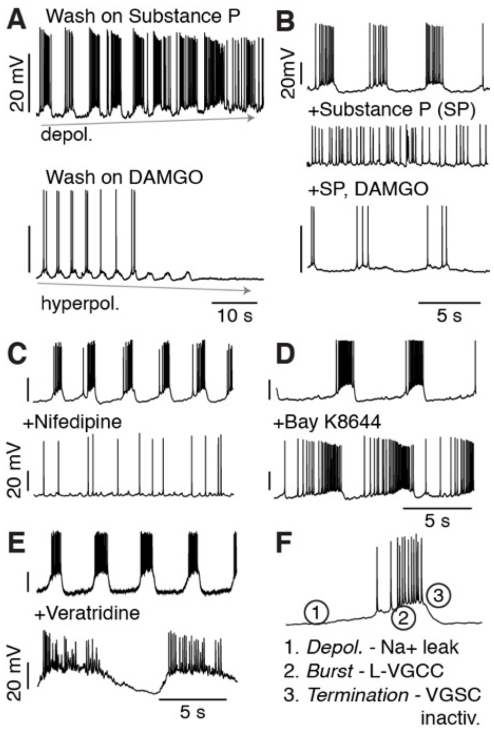


**Fig. S8. The iRO rhythm is voltage dependent.** **A**, iRO rhythm in fast synaptic blockers during bath application of the depolarizing neuromodulator Substance P (200nM, top) or hyperpolarizing neuromodulator DAMGO (200nM, bottom). **B**, Top, baseline. Middle, Substance P. Bottom, Substance P and DAMGO. **C**-**E**, Current clamp recording iRO neurons in fast synaptic blockers at baseline (top) and after bath application of nifedipine (**C**, 2μM), Bay K8644 (**D**, 100nM), and veratridine (**E**, 250nM). Note, baseline rhythm varies from slice to slice, but each neuron was confirmed to be part of the iRO gap junctioned network based on retained rhythmic activity in fast synaptic blockers. **F**, Model for molecular basis of iRO rhythm. $Na^+$ leak channels depolarize the network until VGSC spiking occurs (1) which is converted into a burst by L-type voltage gated calcium channels (2). The burst is terminated by inactivation of VGSCs (3).

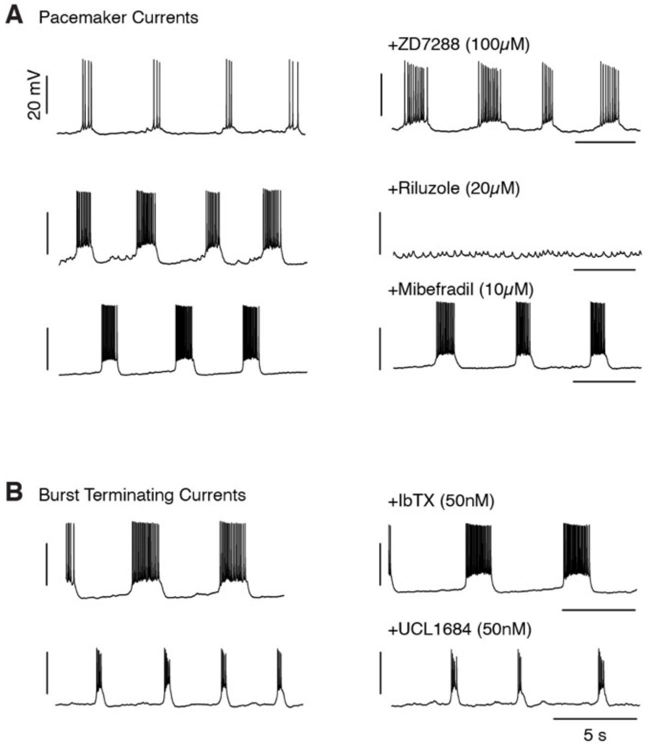


**Fig. S9. Pharmacological analysis of ionic currents required for iRO rhythmic activity. A**, Current clamp recordings of iRO neurons in the presence of fast synaptic blockers. Left, baseline rhythm. Right, rhythm after bath application of pharmacology to antagonize key currents for generic iRO activity. Top, $I_h$ (100µM ZD7288). Middle, persistent sodium (20µM Riluzole). Bottom, T-type calcium channel (10µM Mibefradil). **B**, As in **A,** except pharmacology targeted channels important for burst termination. Top, calcium dependent potassium channels BK (50nM IbTX). Bottom, SK (50nM UCL1684). As in fig. S7, the baseline iRO rhythm varies from slice to slice.

**Table S1. iRO activity in transgenic Cre-induced GCaMP6 lines that label different neural types and compartments of the respiratory neural network.**

| Allele | iRO | Respiratory brainstem compartment | |
|---|---|---|---|
| Vglut2-cre | yes | Glutamatergic | n=4 |
| Gad2-cre | no | Inhibitory | n=4 |
| Vgat-cre | no | Inhibitory | n=3 |
| Glyt2-cre | no | Inhibitory | n=8 |
| RIKEN-Glyt2-cre | no | Inhibitory | n=2 |
| Chat-cre | no | PiCo | n=8 |
| Dbx1-cre | yes | preBötC | n=3 |
| Foxp2-cre | no | preBötC | n=8 |
| Egr2-cre | no | Embryonic parafacial | n=2 |
| Neurod6-cre | no | BötC | n=2 |
| Penk1-cre | yes | Neuropeptide | n=1 |
| Parv-cre | no | BötC | n=1 |
| Tac1-cre | yes | Neuropeptide | n=1 |
| NPY-cre | no | A1/C1 | n=1 |

**Movie S1. The changes in breathing movements related to cries in fig S1**. Neonatal mouse in plethysmograph and ultrasonic vocalization recording chamber. Video shows two breaths that each contain a bisyllabic cry. See fig. S1 for airflow trace and spectrogram. Numbers correspond to the numbers underlying the trace to fig. S1. These are used to align the time in the movie with fig. S1. Note, as the animal inspires the chest/abdomen retract, the head moves forward, and the mouth opens (examples 104-106 and 113-115). During the gap between the two syllables, the same movements are observed (examples 107-109 and 116-118). Thoracic body wall movements trajectories in fig. S1. The body movements and EMG recordings (Fig. 2) are consistent with the airflow oscillation between two syllables being generated by an inspiratory motor program.
